# Functional clustering of splice-altering variants in whole genome sequencing data reveals hidden heritability in rare variant disorder

**DOI:** 10.1101/2023.05.30.542855

**Authors:** Yan Wang, Charlotte van Dijk, Ilia Timpanaro, Paul Hop, Brendan Kenna, Maarten Kooyman, Eleonora Aronica, R. Jeroen Pasterkamp, Leonard van den Berg, Johnathan Cooper-Knock, Project MinE ALS sequencing consortium, NYGC ALS consortium, Jan Veldink, Kevin Kenna

## Abstract

Explaining missing heritability in rare disorders requires effective methods to interpret genetic variants. Sequence-to-function models such as SpliceAI support discovery of splice altering variants but filtering their output to identify pathogenic mutations remains challenging. We developed SpliPath to address 2 unmet needs in this process. First, SpliPath links the output of SpliceAI with reference transcriptomics data. This allows users to identify genetic variants that induce unannotated splice isoforms selectively expressed in disease models or patient tissue. Second, SpliPath aggregates variants with similar functional consequences into collapsed splicing quantitative trait loci (csQTLs) for more powerful genetic association analyses. We first used SpliPath to annotate whole genome sequencing (WGS) of 9,467 ALS patients and controls using RNAseq data from an iPSC model of TDP-43 dysfunction. Through this, SpliPath identified 53 variants predicted to enhance cryptic exon (CE) retention events associated with a core ALS pathomechanism. We then applied SpliPath to 294 ALS patients where both WGS and RNAseq were available and discovered missing genetic risk in the known ALS gene *KIF5A*. This revealed a first of kind intronic mutation hotspot that was validated using minigene reporter assays. Finally, using the same RNAseq data we then predicted 754 candidate csQTL for an independent WGS cohort of 6,625 ALS patients and 2,472 controls. Unbiased genomewide csQTL association testing successfully recovered *KIF5A* and nominated *EPG5* as a potential pathogenic gene. These effects were undetectable using simplistic SpliceAI gene burden tests. Collectively, our study demonstrates the utility of SpliPath for uncovering missing heritability in rare disorders.

## Introduction

Uncovering the missing heritability of rare variant disorders remains a significant challenge. Current methods often cannot capture the full spectrum of genetic contributions, especially in disorders characterized by high genetic heterogeneity and a prevalence of rare variants with low to moderate effect size. This difficulty arises from two primary sources: the complex functional interpretation of genetic variants and the limited statistical power to establish associations between rare variants and phenotypes.

The complexity of functionally interpreting genetic variants arises from the diverse ways in which genetic variants can influence gene function and regulation. Variants can act with incomplete penetrance, and the same genetic variant can have varying effects depending on cellular context and the presence of other genetic factors. In addition, the lack of functional annotations for non-coding regions makes it more difficult to interpret non-coding variants. Aberrant RNA splicing has been increasingly recognized as a key mechanism linking genetic variants to diseases ^1,2^, especially in diseases where affected tissues express diverse splicing isoforms, for example, neurological disorders ^3–5^. Splicing variants can alter splice sites by disrupting splicing regulatory elements, which are genetic sequences bound by spliceosome and splice regulatory proteins. These disruptions can produce dysfunctional transcripts, potentially resulting in disease. Recent advances in deep learning-based prediction frameworks, such as SpliceAI^6^ and MMSplice^7^, have significantly enhanced the identification of splice-altering variants. However, each individual’s genome contains thousands of such variants, whose deleterious effects and relevance to disease remain difficult to assess comprehensively. Additionally, cellular mechanisms protect cells from splicing errors like nonsense-mediated decay can reduce the impact of many deleterious splicing isoforms^8^, complicating the interpretation of their role in disease pathogenesis. These gaps suggest the need for integrated approaches that combine genomic and transcriptomic data to improve variant interpretation.

The other primary challenge in elucidating the genetic basis of rare diseases is the limited statistical power to establish associations between rare variants and phenotypes. Collapsing methods that test the aggregate association between disease risk and multiple rare variants can mitigate the problem of limited statistical power ^9–11^. However, these tests rely heavily on the appropriate selection of “qualifying variants”^12^. The inclusion of benign variants will obscure association signals from *bona fide* disease variants and cause a loss in sensitivity for disease gene discovery. Conversely, excessive stringency in selecting qualifying variants will attenuate power for disease gene discovery by excluding *bona fide* disease variants. Thus, with WGS, effective approaches to enrich true disease variants from the majority of benign variants are needed to enhance the statistical power of rare variant association studies.

To address these challenges, we developed SpliPath, a computational framework to functionally cluster splice-altering variants in WGS dataset into collapsed splicing quantitative trait loci (csQTL), which we define as groups of variants that have similar effects on transcript splicing. In this study, we applied SpliPath to two WGS datasets pertaining to amyotrophic lateral sclerosis (ALS), a complex rare variant disorder with notable unresolved heritability^13,14^. Our work explored the annotation of these datasets using reference transcriptomics datasets that represent distinct categories of disease-relevant contexts. Specifically, we explore the utility of SpliPath in 3 use cases. The first concerns testing whether rare ALS patient mutations could be directly linked to enhancement of disease associated aberrant splicing events reported in RNAseq analyses of a key ALS cell perturbation model. The second involves using paired RNAseq of central nervous system (CNS) tissues from matched patient donors to map hidden heritability in known ALS genes. Finally, the third explores using unpaired RNAseq of tissue from unmatched donors to guide unbiased disease gene discovery in a genomewide case-control association testing setting. Through this we showcase the functionalities of SpliPath and demonstrate how it can be deployed to refine the search space for pathogenic variants in challenging rare disorders.

## Results

### SpliPath uses disease relevant transcriptomics data to annotate splice-altering variants in patient WGS

SpliPath is designed to identify and functionally cluster splice-altering variants in WGS that have similar effects on RNA splicing. We refer to these variants as csQTL. The first step of this process is to link genetic variants observed in patients with reference splicing junctions that are detected in transcriptomic analyses of patients or model systems (Figure 1A). To achieve this, we require “matching” of the exact splicing junction detected from splicing analyses with the expected splice site gain or loss events predicted by SpliceAI. Here, a splicing junction refers to the combination of a donor and an acceptor splice site where an intervening intronic sequence has been removed to connect two exons. For each genetic variant, SpliceAI can predict changes in splicing activity at up to 4 possible splice site locations. These 4 predictions correspond to distinct donor site gain, donor site loss, acceptor site gain and acceptor site loss events. When an observed splice junction matches fully with a set of SpliceAI predictions then SpliPath classifies the pairing as a “full match” event. SpliPath can also report “partial match” events, which correspond to pairings where splice site usage at an observed junction only partially corresponds with a set of SpliceAI predictions (Figure 1B, S1). SpliPath matching can performed in the context of both paired and unpaired WGS and transcriptomic datasets (Figure S2). Variants assigned to csQTL can be modelled in aggregate to assist with resolving variants of unknown significance or to give power to unbiased genomewide discovery analyses.

**Figure 1.**
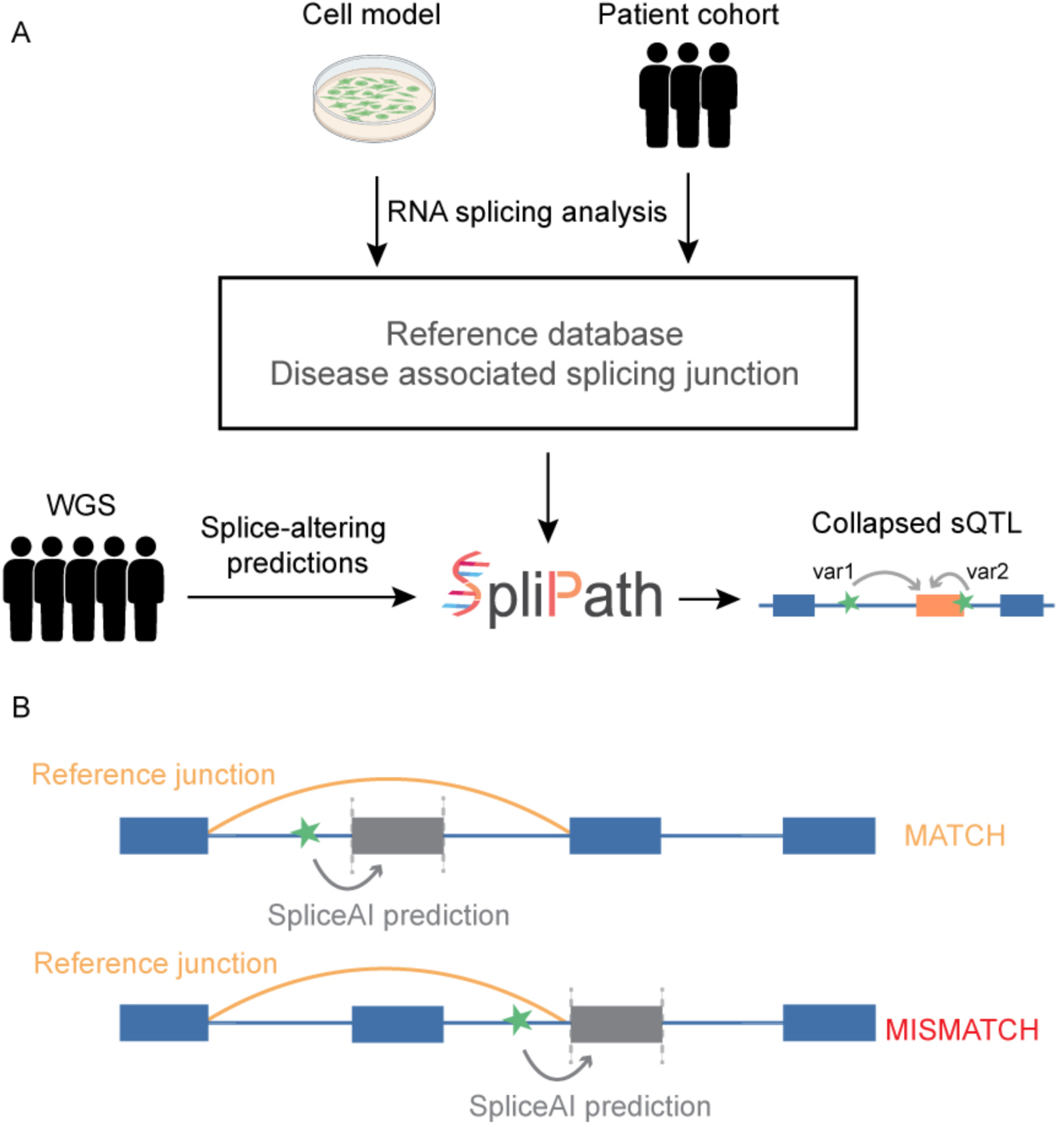
**SpliPath to match WGS mutations and RNAseq derived reference junctions for csQTL analyses**. A) SpliPath takes WGS data and RNAseq derived splice junction maps as input. Splice junction maps can be used to create reference databases using different strategies. In this study we nominate junctions of interest first by applying differential splicing analyses to RNAseq of ALS cell perturbation models and second by applying outlier analyses to RNA-seq of ALS patient tissue. Variants that are assigned to similar splicing phenotypes are aggregated as csQTL for downstream genetic association analyses. B) SpliPath first establishes candidate pairings between WGS derived genetic variants and RNAseq derived splice junctions by positional matching. These initial matches are then further filtered by using the output of SpliceAI and RNAseq analyses to distinguish “match” from “mismatch” pairings (see Methods, further details in Figure S1). Grey dashed lines represent splice site losses predicted from SpliceAI annotation of genetic variants (green stars). Golden arcs represent splice junction observed in the reference RNAseq datasets. The splice junction is the combination of a donor and an acceptor splice site where two exons are connected. In panel B, the reference splice junction suggest a skipping of the second exon.

### Use of SpliPath to annotate ALS WGS for csQTL associating with cryptic exons induced in cellular disease model of ALS pathology

The depletion of nuclear TDP-43 is a neuropathological hallmark of ALS and was reported to induce cryptic exon (CE) inclusion in 179 genes in iPSC derived neurons ^15^. Two of these CE, found in *STMN2* and *UNC13A*, are already known to be neurotoxic. Importantly, it has also been demonstrated that the CE inclusion event within *UNC13A* is enhanced by cis-acting genetic variants that associate with ALS risk and clinical phenotype in GWAS ^15,16^. An open and high priority research question for the field is whether additional undiscovered genetic variants also regulate other toxic CE induced by TDP-43 dysfunction. As a first step towards addressing this question, we used CE profiles from Brown et al^15^ to build a reference database of target splice junctions. As a pre-processing step we filtered the Brown dataset for CE that we could also detect in RNAseq of ALS postmortem tissue. We then used our SpliPath workflow to annotate 2 ALS WGS datasets for candidate csQTL which are predicted to enhance TDP-43 associated CE retention events. These WGS datasets included 294 ALS patients and 76 controls from the NYGC ALS sequencing consortium and an independent set of 6,625 ALS patients and 2,472 controls analysed by the Project MinE consortium. For the NYGC cohort, paired RNAseq of postmortem brain and spinal cord material from matched donors was also available. This paired data enabled further comparisons of individual genotype and splicing phenotypes within individual subjects.

Collectively, our analyses revealed 53 candidate csQTLs across the 2 cohorts (Table S1, S2). The overall frequency of these csQTL was marginally higher in patients than controls, though this difference did not reach statistical significance (Pooled analysis of csQTL with MAF<1%: ALS MAF=0.68%, control MAF=0.59%, P=0.54. Pooled analysis of csQTL with MAF<0.1%: ALS MAF=0.34%, control MAF=0.27%, P=0.57). Our set of nominated csQTL did not recover the 2 known CE modifier variants in *UNC13A* (rs12973192, rs12608932) ^15,16^. These variants were assigned SpliceAI prediction scores below the standard confidence threshold of 0.2 (rs12973192=0.02; rs12608932=0.01), highlighting a noteworthy limitation for the current generation of AI-based splice prediction tools that could be improved in future releases. All the junctions associated with the 53 nominated csQTL were detectable in patient tissue, however in bulk tissue most cells do not exhibit marked TDP-43 loss of function and CE expression levels are too low for detailed analyses. For example, ALS associated variants in *UNC13A* provide a major enhancement of CE retention in minigene reporter assays when concomitant TDP-43 knockdown is performed^15^. However, in comparisons of CE read counts across carriers and non-carriers of the *UNC13A* SNPs we observed only a very subtle shift in global *UNC13A*-CE levels (Figure S3A, B). Accordingly, the paired WGS and RNAseq dataset from the NYGC provides limited power to test whether rarer csQTL enhance CE retention in a manner that requires concomitant TDP-43 dysfunction. However, the dataset could be powered to detect csQTL that are strong enough to overcome TDP-43 repression and induce a more constitutive CE retention. Amongst 13 candidate csQTL carried by patients in the NYGC dataset, we observed that these criteria appeared to be fulfilled by a chr12:88086638A>T variant predicted to induce CE retention within the *CEP290* gene. The carrier of this variant exhibited a statistically significant upregulation of the corresponding junction using LeafCutterMD outlier analyses (P=2.18×10^-5^, Figure S2, S3C, D). This particular variant exhibited similar frequencies in patients vs controls (ALS MAF=0.16%, control MAF=0.16%, P = 0.64) and no clear association with patient survival time (P = 0.96, HR = 1.01, CI = 0.45-1.58, Cox proportional hazards regression). However, the observation demonstrates that SpliPath can successfully bridge rare WGS variants with disease relevant perturbation associated splice disruptions observed in independent cell models. As such our database of csQTL assignments between ALS patient variants and TDP-43 CE provides a useful resource for future targeted research into genetic regulation of TDP-43 splicing defects in ALS.

### Use of SpliPath to annotate ALS WGS for csQTL associating with unannotated splice junctions in paired RNAseq

Next, we sought to nominate candidate disease splice junctions through statistical outlier testing analyses of RNAseq data from the NYGC ALS sequencing consortium. The dataset includes a total of 1,044 RNAseq profiles, each representing 1 of up to 4 brain or spinal cord regions within a cohort of 370 unique individuals (294 cases, 76 controls). Using LeafCutterMD^17^ and our splice junction classification procedure (see Methods), we detected a total of 509,144 unique unannotated splice junctions where P values from outlier testing exceeded a nominal significance threshold. We observed no significant difference in the frequency of outlying splice junction in patients vs controls for any of the 4 tissues (frontal cortex: p=0.81; motor cortex: p=0.24; cervical cord: p=0.08; lumbar cord: p=0.47; T test, Figure 2). Using SpliPath, we then reduced this set of 509,144 junctions to 754 candidate csQTL derived from the patient samples (Table S3, Figure S4. Results from csQTL analyses of control samples also provided Table S3). These junctions assigned to candidate csQTL thus 1) were unannotated, 2) met a nominal statistical threshold for classification as outlying in at least one subject, and 3) were fully or partially matched with the predicted consequences of genetic variants that were carried by the same subjects exhibiting outlying junction usage. As the primary focus of our research is the annotation of rare variants, these events were also restricted to junctions where the matched genetic variant exhibited a miner allele frequency below 0.1% or homozygous carrier frequency below 0.1% among gnomAD (v3.1.2)^18^ non-neurological disease controls.

**Figure 2.**
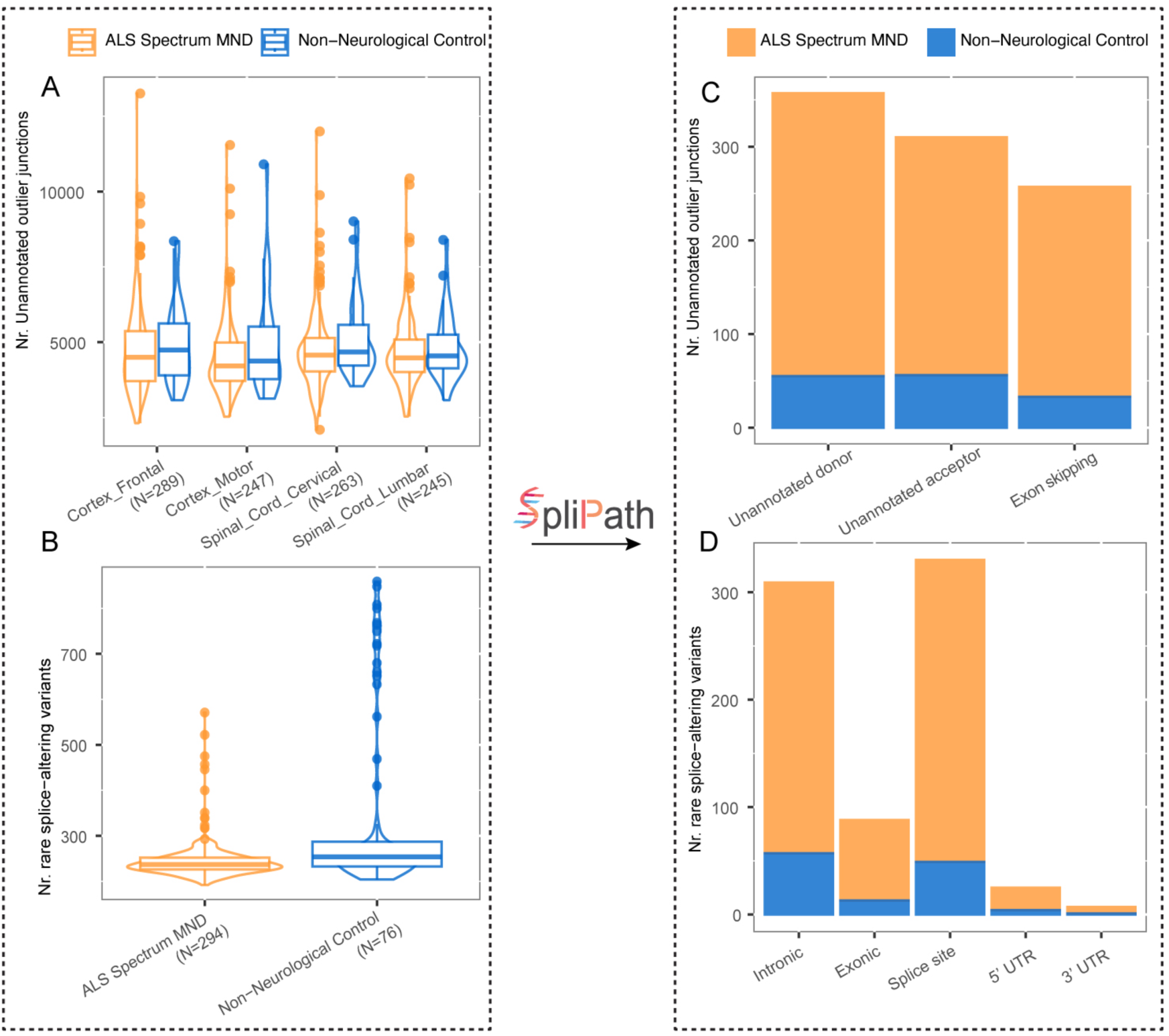
**SpliPath prioritized disease-relevant splicing junctions by annotating csQTL using paired WGS and RNAseq dataset.** A) Number of unannotated outlier splice junctions per individual by LeafCutterMD analysis of 1,044 RNAseq samples of brain and spinal cord from 370 individuals in NYGC ALS cohort. B) Number of rare splice-altering variants predicted by SpliceAI for patients and controls. From them, SpliPath nominated csQTL that involved C) 754 unique unannotated outlier junctions and D) 611 unique observed splice-altering variants in patients’ tissues (csQTLs nominated from both patient and control samples are provided in Table S3).

Before using the 754 candidate csQTL to annotate WGS for additional patients in the Project MinE dataset, we first conduct a more detailed assessment of csQTL impacting known ALS genes within the paired WGS and RNAseq datasets from the NYGC consortium. Within only 76 controls, the NYGC cohort was underpowered for csQTL association testing. However, nonetheless could use the paired datasets to establish 2 clear likely pathogenic variants (Figure 3). The first variant disrupted a consensus splice site region in the *TBK1* gene (chr12:64496407G>C, SpliceAI score = 1.0, n=1 patient). The second variant was 58bp away from the intron/exon boundary of exon 27 in *KIF5A* gene (chr12:57582544A>C, SpliceAI score = 0.64, n=1 patient). Using BPP^19^, we predicted that the *KIF5A* variant corresponded to a branchpoint disruption. The splice sites loss of *TBK1* exon 16 and *KIF5A* exon 27 predicted by SpliceAI exactly matches the exon skipping events that we observed in RNA-seq data of the two variant carriers (Figure 3B, 3C).

**Figure 3.**
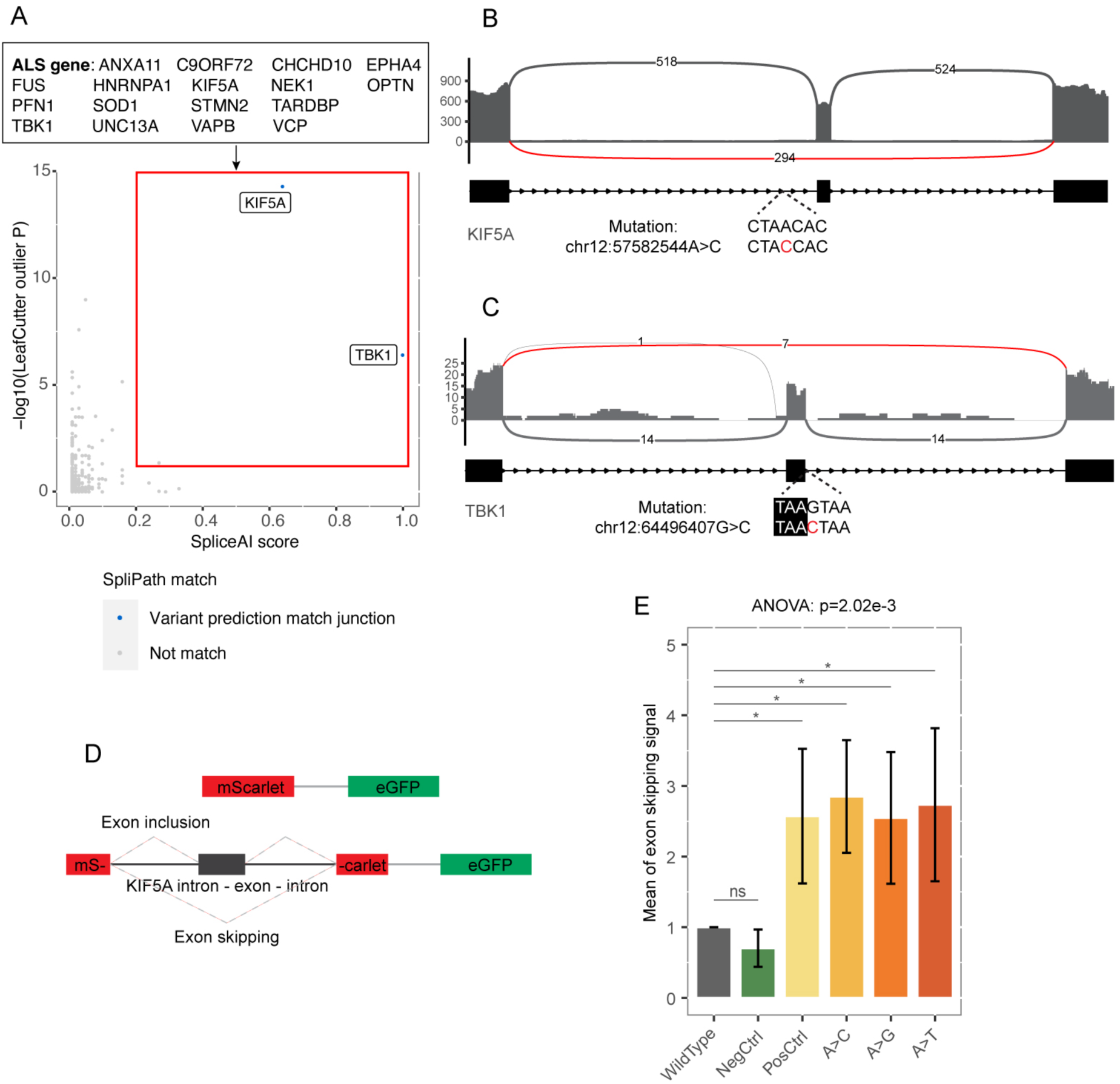
**SpliPath annotation of csQTL in paired WGS and RNAseq dataset reveals pathogenic variant in *KIF5A* and *TBK1*.** A) Each dot in the scatter plot shows a putative csQTL: a rare variant within the csQTL search space (Method) of an unannotated junction in a patient motor cortex sample. The SpliceAI score of the rare variant is shown in x axis and LeafCutter outlier P value of the unannotated junction is shown in Y axis. The red box shows the initial thresholds for csQTL inclusion: LeafCutter outlier P value < 0.05 and SpliceAI score ≥ 0.2. By matching the SpliceAI prediction of variants with the unannotated junctions, SpliPath identified csQTL (blue) in *KIF5A* and *TBK1* among putative hits (grey) in 17 known ALS genes. The csQTLs are intronic splice-altering variants causing unannotated exon skipping events in patients’ RNAseq profile in B) *KIF5A* and C) *TBK1*. D) Design of minigene reporter to validate the exon skipping effects of *KIF5A* variants discovered in DNAseq only data. The plasmid contained an eGFP and a split mScarlet in which the *KIF5A* intron 26 - exon 27 - intron 27 sequences with(out) variants were inserted (See design of whole plasmid in Figure S7) E), Bar plot of exon skipping signals of minigene reporter assays. The negative control variant was from gnomAD (v3.1.2)^18^ database with allele count = 5 in non-neurological disease group, and the positive control variant was a splice consensus region variant that we reported previously^11^ (Table. S1). Quantification of exon skipping was based on normalized median mScarlet signal over 3 technical replicates and the experiment was repeated 4 times. There were no differences of exon skipping signals between cells with wildtype *KIF5A* sequence insertions and negative control sequences insertions (“ns”), while exon skipping signals of cells with positive control and target sequences insertions were significantly higher than that of cells with wildtype *KIF5A* sequence insertions (“*”, P.adj < 0.05, T test, FDR adjusted; error bars represent mean ± standard deviation).

The *KIF5A* variant was an especially interesting observation as this mutation had not been reported in prior genetic analyses of the same patient cohorts ^11^. Further analyses also revealed that the mutation occurred deeper into intronic sequences than is detectable when using the precomputed scores released for by the authors of SpliceAI^11^. This latter point was surmountable in our study because our prior preselection of unannotated patient junctions from RNAseq data enables more detailed screening of local variant search spaces. Evaluation of the mutated position in the Project MinE datasets revealed 2 additional carriers of variants at the same location (hg19, chr12:57976327A>C n=1 patient and chr12:57976327A>G n=1 patient). To further substantiate that this set of variants indeed induce the predicted exon 27-skipping event, we designed a minigene reporter assay based on a construct used in a recent massively parallel splice reporter assay^20^ (Figure 3D). This construct included an mScarlet reporter gene where the coding sequence is disrupted by insertion of *KIF5A* exon 27 with the entirety of the flanking introns. Insertion of mutations that induce exon skipping into this construct are accordingly expected to provide a detectable rescue of fluorescent intensity. Our assay revealed no differences in exon skipping signals between cells expressing the wildtype *KIF5A* sequence insertions and negative control DNA variants. However, exon skipping signals were significantly higher for both positive control variants that we identified at the intron-exon boundaries in previous work^11^, and for the newly discovered intronic variants that we report in this study (Figure 3E). Combined these observations reveal a first of kind intronic mutation hotspot for ALS that can be evaluated in future genetic testing of ALS patients without the need for parallel RNA-seq analyses.

### Use of SpliPath to annotate ALS WGS for csQTL associating with unannotated splice junctions in unpaired RNAseq

As a final use case, we evaluated the utility of SpliPath to support disease gene discovery in large case control association analyses where WGS data is annotated using csQTL derived from transcriptomics analyses of an independent study cohort. For this, we used the 754 candidate csQTL junctions described in the previous section to annotate WGS of 6,625 ALS patients and 2,472 controls from the Project MinE cohort. Our analyses revealed 479 single nucleotide variants (SNVs, carrier frequency < 0.1%) that could be assigned to one of the reference junctions (Table S4). In contrast, a total of 124,724 rare SNV achieved a precomputed SpliceAI scores ≥ 0.2 in the same dataset. The inclusion of SpliPath based reference transcriptomics matching thus had a dramatic effect on the size of the target search space. We then used firth logistic regression to test each csQTL for association with ALS risk. For comparison, we also performed gene burden analyses that were restricted to all variants with precomputed SpliceAI scores ≥ 0.2 (Figure 4, Table S5). By incorporating the magnitude of LeafCutterMD analyses, the *KIF5A* exon 27 skipping csQTL is readily discoverable as a lead candidate for further validation in SpliPath analyses (Figure 4, Table 1). In total, this csQTL represents an aggregation of 6 independent variants that are observed within 8 patients and 0 controls (Table S6). Although, this data is in line with a high odds ratio known to associate with exon 27 skipping^11^, larger sample sizes are required to demonstrate a significant P value. Conversely, the *KIF5A* association is in no way observable when a simple SpliceAI based gene burden analysis is applied (OR=0.94, p=0.91. Fig 4).

**Figure 4.**
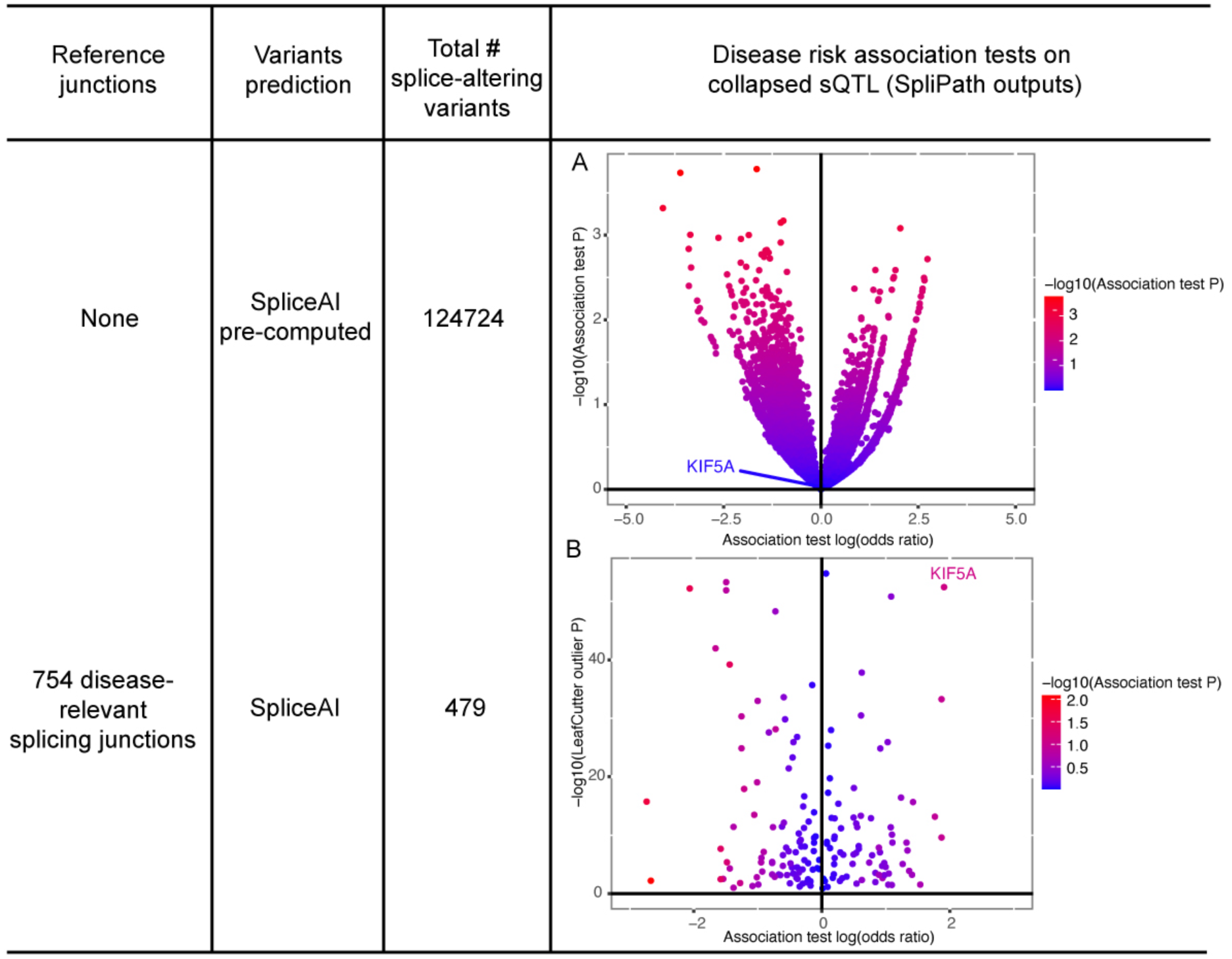
**csQTL association tests can support disease gene discovery in genomewide analyses.** Association tests are performed for A) gene level aggregation of all splice-altering variants nominated by SpliceAI without the use of reference transcriptomic information and B) csQTL mapped to 754 disease-relevant splicing junctions using SpliPath. In panel A, association tests are performed for 124,724 splice-altering variants annotated as having precomputed SpliceAI scores ≥ 0.2. Each dot shows the association of splice-altering variants withih a gene with disease phenotype in Project MinE WGS. In panel B, there are 479 csQTL SNVs mapped to the 754 splicing junctions. Each dot shows a splicing junction in NYGC dataset with an outlier splicing P value (Y axis) and the association test odds ratio on log scale (X axis) of the csQTL in Project MinE WGS. Dots are colored by the association test P value. Association analyses are restricted to SNV and testing genes/csQTL that have at least 3 observations in Project MinE.

**Table1:** Top candidate csQTL from SpliPath annotation of Project MinE WGS cohort.

| Gene | Event | Junction | Analysis | # Variants | Case carriers | Control carriers | P |
| --- | --- | --- | --- | --- | --- | --- | --- |
| KIF5A | Exon skipping | 12:57581952:57583100:+ | allelic, SNV only | 6 | 8 | 0 | 0.0804 |
| KLHDC9 | Exon skipping | 1:161099062:161100060:+ | allelic, SNV + INDEL | 5 | 8 | 0 | 0.0713 |
| KLHDC9 | Exon skipping | 1:161099062:161100060:+ | allelic, SNV only | 4 | 7 | 0 | 0.1225 |
| EPG5 | Unannotated donor site | 18:45901167:45903939:- | recessive, SNV + INDEL | 1 | 6 | 0 | 0.2259 |
| FAM102B | Unannotated acceptor site | 1:108629657:108635430:+ | recessive, SNV + INDEL | 1 | 6 | 0 | 0.1773 |
| SLC25A46 | Unannotated donor site | 5:110748678:110755464:+ | allelic, SNV + INDEL | 2 | 5 | 0 | 0.2883 |
| NPRL3 | Unannotated donor site | 16:88890:89764:- | allelic, SNV + INDEL | 2 | 5 | 0 | 0.2428 |
| EDIL3 | Exon skipping | 5:84180521:84384307:- | allelic, SNV + INDEL | 1 | 5 | 0 | 0.4871 |
| NINJ1 | Unannotated donor site | 9:93125062:93125656:- | allelic, SNV + INDEL | 1 | 5 | 0 | 0.1065 |
| NINJ1 | Unannotated acceptor site | 9:93125749:93126409:- | allelic, SNV + INDEL | 1 | 5 | 0 | 0.1065 |

\* Top events include csQTL observed in 5 or more patients but 0 controls. A complete table of csQTL with case control frequencies is available in Table S5.

Finally, following the inclusion of indels and recessive disease models, we observed 7 additional genes where the set of csQTL linked variants were both absent from controls and present in ≥ 5 patients. Larger sample sizes will be required to robustly evaluate statistical associations for these csQTL. However, a notable example amongst these observations is *EPG5*. Within *EPG5* we observed 6 homozygous carriers of a chr18:g.45903935T>TTCAC variant. This gene is associated with Vici Syndrome, a condition exhibits early-onset CNS developmental defects and abnormalities in immune system, heart and skin. Interestingly, previous studies on *EPG5* knockout mouse models showed key ALS characteristics, including neuron loss, TDP-43 aggregation in motor neurons, and muscle atrophy^21–23^. The csQTL we observed causes elongation of exon 25 and is predicted to introduce an insertion of 11 amino acids in protein. We observed a significant correlation between expression of this junction and mutation dosage across the 13 heterozygotes, 356 non-carriers and 1 homozygous variant carrier in the NYGC cohort (P< 2×10^-16^ in all the 4 tissues datasets, Figure S4). In the Project MinE dataset, the homozygous variant was found in 6 patient carriers and 0 control carriers, and Hardy Weinberg Equilibrium tests suggested a statistically significant over-representation of homozygous variants among patients (P=3.2×10^-4^).

## Discussion

Studies of simple Mendelian disorders can leverage familial segregation patterns and known disease gene panels to conduct extensive pre-filtering for causal splice-altering mutations ^2,24^. However, the genetic architecture for many diseases, for example ALS, is more complex. Although disease risk is dominated by rare variants, very few patients exhibit evidence for Mendelian inheritance and most heritability remains unexplained. The capacity to detect carriers of pathogenic intronic variants will also become increasingly urgent when it is needed to determine the eligibility of patients for emerging gene targeting antisense oligonucleotide therapies such as Jacifusen^25^ or Tofersen^26^. We therefore use SpliPath to nominate disease-associated splicing events from publicly available RNAseq datasets and establish csQTL for WGS annotation and association mapping. Importantly our framework supports the use of both paired and unpaired transcriptomcis datasets. This is important for disorders like ALS, as key disease tissues are not accessible for transcriptomic analyses in living patients. However, this study design can readily be reused for use cases beyond ALS, most especially for disorders with a challenging rare variant genetic architecture and where disease relevant transcriptomics datasets are already publicly available.

The first question we explored in our work was whether SpliPath could be used to link individual genetic variants with splicing defects induced by disease relevant perturbations in cell models. This was a vastly different approach to traditional sQTL discovery strategies, where paired genetic and transcriptomic data for a reference cohort are used to identify sQTL based on statistical association between genotype and intermediate molecular phenotypes. For these analyses we focused on leveraging TDP-43 depletion experiments in ALS cellular models. We selected this system as the experiments have had a very high impact on ALS research and model a major neuropathological process that can be detected in patient tissues. Furthermore, the prior discovery of *UNC13A* CE modifier variants raised the question of whether additional genetic variants also regulate toxic CE retention events in patients. Using SpliPath, we nominated 53 csQTLs for TDP43-dependent CEs that could be assigned to genetic variants carried by ALS patients. Not all of these variants will necessary prove pathogenic, as suggested by our characterization of the *CEP290* csQTL. However, the csQTL we identified provide a logical focus point for more extensive testing of CE modification under conditions of TDP-43 dysfunction. One strategy to evaluate the CE enhancement by our csQTL in a high throughput manner would be to adapt massively parallel reporter assay systems that have previously been deployed to investigate thousands of candidate splice altering variants^20^. In this context variant effects would be tested with and without TDP-43 knockdown. Alternatively specific CE variants could be prioritized for more detailed investigation in a low throughput manner. Genes with independent biological evidence for ALS association or higher patient frequencies could be introduced into reference iPSC lines using CRISPR editing. This could enable well controlled investigations into the effects of csQTL on endogenous transcripts in iPSC derived motor neurons, with the potential to then pursue more detailed analyses into whether and how csQTL modifiers might impact relevant cell phenotypes.

Our analyses of splice junction profiles in brain and spinal cord revealed thousands of unannotated splice junctions in each subject. Notably, we observed similar overall frequencies of these junctions amongst the ALS and control groups, suggesting that most events detected in patients are likely non-pathogenic. Likewise, our screening of DNA variants predicted to be splice disrupting using SpliceAI, revealed hundreds of candidate variants per individual. Collectively, these observations highlight the challenges of identifying diseases relevant splice altering variants in unimodal analyses. We therefore first explored the capacity of SpliPath to mitigate these challenges by combining data across modalities using paired DNA and RNA sequencing datasets. Through this, we identified novel ALS variants that induced *TBK1* exon 16 skipping and *KIF5A* exon 27 skipping respectively. The novel variant identified in *TBK1* impacted a consensus splice site and could thus be discovered through conventional DNAseq only analyses. At the protein level, the *TBK1* exon 16 skipping event is expected to result in deletion of functional coiled-coiled domain 2^27^, in-keeping with proposed loss of function models reported in previous studies^28,29^. However, the novel *KIF5A* variant was located 58bp upstream of exon 27 and was not readily identifiable in standard DNAseq only analyses. Additional variants impacting this position were identified in 2 Project MinE patients. We validated the exon-skipping potential of all SNVs at this deep intronic position using a minigene reporter assay, substantiating that this nucleotide is key in regulating exon 27 inclusion. We have demonstrated the selective pathogenicity of exon 27 skipping in previous work^11^. However, the intronic variants we describe here were not recognizable in prior studies and the current study represents the first discovery of an ALS associated intronic mutation hotspot. This finding further substantiates that future clinical genetic testing of ALS patients needs to look beyond protein coding mutations and known repeat expansions.

We also used the NYGC splice junctions to annotate an independent unpaired WGS dataset that included 6,625 patients 2,472 controls from Project MinE. Similar to our CE annotation analyses, this demonstrates how publicly available RNAseq datasets can be used to annotate rare variants in new WGS data. This work also showcases the benefits of our functional clustering strategy, where 6 exon 27 skipping linked variants of *KIF5A* are aggregated into a single csQTL for analysis. This step supported the ready prioritization of *KIF5A* exon 27 skipping as a lead candidate in unbiased gene discovery analyses (OR=6.74), a result that could not be achieved by simply using SpliceAI scores as a filter for gene burden analyses (OR=0.94). More generally our SpliPath approach transformed a search space of 124,724 splice-altering variants nominated through basic SpliceAI annotation of Project MinE WGS, to 479 csQTLs which were assigned to unannotated splice junction consequences readily observable in a large reference panel of ALS patient RNAseq profiles. Our work also nominates homozygosity for an exon 25-extending variant of *EPG5* as a potential risk factor for ALS. In our analysis, the association between variant genotype and junction expression level has been validated by paired RNAseq and WGS data. We also observed that the homozygous genotype is indeed overrepresented in the Project MinE patient group. Previous studies on *EPG5* knockout mouse models showed key ALS characteristics, including neuron loss, TDP-43 aggregation in motor neurons, and muscle atrophy^21,22^. Human *EPG5* has been associated with Vici Syndrome, a condition exhibits early-onset CNS developmental defects, where patients carried splicing and missense variants that truncate the protein^30^. The patients carried the homozygous variant in our study, however, presented phenotypes different from Vici Syndrome, which could be due to the variant having a distinct effect on protein function compared to those seen in Vici Syndrome. Larger cohorts will be required to further test the association of *EPG5* with ALS, but for now the observation serves to highlight the value of also considering recessive models in our csQTL framework.

One limitation of our study is that our sensitivity to detect disease-relevant splicing junctions was constrained by the technical noise and lack of single cell resolution that is inherent to short read RNAseq of bulk tissues. Future datasets that increase the quality of reference transcriptomics through single cell resolution, long read sequencing or larger sample numbers will create new opportunities to discover missing ALS splicing junctions. In theory this could be particularly advantageous for scenarios such as TDP-43 depletion, where only a small subset of cells express key disease junctions in patient tissues. Naturally, large WGS datasets will also provide greater power to validate *bona fide* pathogenic variants in genomewide csQTL analyses. Finally, our study incorporated SpliceAI for the prediction of splice-altering variants, but sequence to function models for splice variant prediction are still under active development and future advances from tools such as Pangolin^31^ may allow for new performance gains. The potential gains of this are again underscored by the fact that certain key splice variants, such as the CE enhancing variants of *UNC13A*, are not well catered for using SpliceAI.

In conclusion, we developed SpliPath to functionally cluster splice-altering variants into csQTL for WGS analysis. By applying SpliPath to a selection of ALS datasets, we demonstrate that transcriptomics data from cell models, paired RNAseq of patient tissue and unpaired RNAseq of patient tissue can all be leveraged to add informative annotations to rare variants identified by WGS. We also demonstrate that csQTL analyses can provide advantages over analyzing rare splice mutations individually or in under less refined whole gene collapsing analyses. Our work provides both novel resources and insights for ALS research but also describes how SpliPath can readily be reused in any disorders where rare genetic variants contribute to disease heritability and relevant reference transcriptomics datasets are available. Notably our work also highlights a functional clustering strategy which can also be extended beyond splicing to equivalent sequence dependent molecular phenotypes such as promoter function or polyadenylation.

## Method

### RNAseq and DNAseq datasets

This study used WGS from 2 ALS cohorts (Project MinE, NYGC ALS sequencing consortium) and RNaseq from 2 ALS relevant datasets (i3Neuron models of ALS related TDP-43 dysfunction^15^ and multi-tissue RNAseq of ALS patients by the NYGC ALS sequencing consortium). The Project MinE data includes WGS for 6,652 ALS patients and 2,472 controls. Full details of this cohort have been reported in van Rheenen al^13,14^. Data from the NYGC ALS sequencing consortium includes full WGS for 294 ALS / motor neuron disease (MND) spectrum patients and 76 neurotypical controls, as well as paired RNAseq of up to 4 brain and spinal cord regions sampled from the same matched donor cohort (motor cortex, frontal cortex, cervical spinal cord, lumbar spinal cord). RNAseq data for TDP-43 knockdown/control i3Neurons were retrieved from the European Nucleotide Archive (PRJEB42763) ^15^. For a secondary check for ALS related cryptic exons, we used RNAseq data for TDP-43 negative/positive neuronal nuclei from postmortem patient tissue, retrieved from the Gene Expression Omnibus (GSE126543)^32^.

### Alignment and pre-processing of RNAseq and DNAseq datasets

All RNA-seq data was aligned to the GRCh38 reference genome using STAR v2.7.3a^33^. All transcript annotations were based on Ensembl version 98. To enhance capture of unannotated junctions in downstream analyses, STAR was used in two-pass mode (whereby unannotated junctions identified during first pass mapping are used to generate a refined splice junction annotation file for second pass mapping). All WGS data from the NYGC was aligned to the hg38 reference genome using BWA^34^. Variant calling and individual genotyping were performed in accordance with best practices outlined for the Genome Analysis Toolkit (GATK v4.1.0)^35^. Processing of the WGS dataset from Project MinE is described by van Rheenen et al^13^. 27 sample duplication events between Project MinE and the NYGC dataset were identified by using KING to analyze genetic relatedness^36^. These relatedness analyses were conducted using a quality controlled set of common variant markers that were extracted from VCF files using plink v1.9^37^. Quality control filters for the common variant markers included filtering for a set of biallelic LD pruned single nucleotide variants that exhibited a MAF>0.01, genotype call rate>0.9 and no evidence for deviation from Hardy Weinberg equilibrium (P<0.001).

### Outlier splice junction identification

RegTools v0.5.2^38^ was used to define candidate splice junctions based on the detection of split reads within the aligned RNAseq bam files. Split reads were only considered in the event that they mapped to the reference genome with a minimum overlap of 6bp at both ends. LeafCutterMD^17,39^ analysis was used to detect outlier splicing junctions of each tissue. This involves two steps: 1) LeafCutter intron clustering method was performed on the splicing junction files, with parameters: minimum 10 reads in a cluster, minimum 0.0001% of reads in a cluster that support a junction, and maximum intron length of 500k bp; 2) LeafCutterMD analysis on the clustered introns with parameters: maximum 500 introns in a cluster and minimum 10 reads mapped to a cluster. The junctions with an outlier P value < 0.05 were considered as outlier junctions. The outlier splicing junctions were classified as being “annotated” or “unannotated” by comparison to reference splice junction coordinates retrieved from Ensembl 98 and the Snaptron database^40^. Unannotated splice junctions were classified as being either unannotated exon skipping junction, unannotated donor junction, or unannotated acceptor junction. For designation as an unannotated exon skipping, we required that the data supported skipping of annotated exons without creating unannotated donor/acceptor sites. For designation as an unannotated donor/acceptor junction, we required that an observed unannotated donor/acceptor site was paired with an annotated splice site. More complex splice aberrations involving multiple genes or multiple unannotated splice sites were regarded as low confidence events and excluded from our analyses in this study. Candidate junctions occurring within ENCODE blacklist regions and sex chromosomes were also excluded ^41^. To obtain disease-relevant splice junctions of higher confidence, we required that the outlier junctions from analysis of NYGC RNAseq dataset were reproducible across RNAseq profiles of at least 2 tissue types from the same donor. For each outlier junction, Fisher’s method in *poolr* package was used to combine the tests of multiple tissue types.

### Obtaining reference junctions from TDP43-dependent cryptic exons

The differential splicing analysis of TDP43-KD i^3^Neurons and controls were retrieved from Brown et al‘s work ^15^. The corresponding splicing junctions of the cryptic exons were those labeled as cryptic and with FDR < 0.05. For these junctions, we examined their expression in RNAseq profiles of postmortem brain and spinal cord in NYGC cohort. The reference junctions for csQTL analysis were those that were detectable (read count >= 1) in any patient’s tissue but not detectable in tissue of more than 7 (10%) controls.

### Prediction of splice-altering consequences for DNA variants

We annotated DNA variants with predictions scores from SpliceAI v1.3^6^. We scored the predicted effect of every observed minor allele on every potential splice site located within 500bp of the variant position. The final SpliceAI score was taken as the maximum delta score predicted for any given donor gain, donor loss, acceptor gain, and acceptor loss event. DNA variants were only considered as potentially splice-altering in the event of a SpliceAI score ≥ 0.2. For DNA variants in Project MinE in build hg19, the variants were first lifted to hg38 using CrossMap^42^ and then annotated using SpliceAI.

### Matching splice-altering variants with reference junctions

For any given reference junction, we restricted our DNA variant search space to include the entirety of the disrupted intron plus both flanking exons. SpliceAI generates four predictions for the effects of a genetic variant: donor site gain, donor site loss, acceptor site gain, and acceptor site loss. The criteria for ‘matching’ a SpliceAI prediction with a reference junction are as follows: 1) If SpliceAI predicts the gain of an unannotated donor/acceptor site, a match is considered when a junction uses this unannotated site; 2) If SpliceAI predicts the loss of an annotated donor/acceptor site, a match is considered when a junction omits this annotated site while using another annotated or unannotated site. Predictions that exceed a delta score threshold of 0.2 were used for matching with reference splicing junctions. In most instances, multiple SpliceAI predictions can pass the threshold for a single genetic variant. Since SpliceAI is able to predict the changes of up to four donor and acceptor sites, it is possible that a splice junction is not align with all of them. We classify a splice junction as a “partial match” if it aligns with some of a variant’s predictions, and as a “full match” if it aligns with all predictions (detailed Figure S2). The splice-altering variants that fully or partially matched to the same reference junction were grouped as a csQTL.

### Case-control association testing

Case-control association tests were performed using the Rare Variant Analysis Toolkit (RVAT) ^43^ (https://github.com/KennaLab/rvat). Firth penalized logistic regression was used to model case-control status with respect to rare variants burden while correcting for sex, sequencing platform and the first 4 principal components derived from PCA of common variant profiles. Rare variants were only included for association testing if carrier frequency within the cohort was below 0.1%^9,10^, under either an allelic or recessive genetic model. Genomewide rare variant association screens were performed under both allelic and recessive model.

### Survival analysis

Survival analysis was performed using proportional hazards regression model (coxph function in R survival package v3.5-5), correcting for sex and the first 4 principal components derived from PCA of common variant profiles as well.

### SpliPath package

The SpliPath R package was developed to support reuse of our workflow for any desired combination of input WGS and RNAseq datasets. The SpliPath package also includes shiny browsers to visualize and manually inspect candidate csQTL in detail. We have developed one browser tailored to paired DNA-RNAseq datasets, where RNAseq and WGS data are both derived from the same donor (example browser: https://yanwang271.shinyapps.io/splipath_browser/). We have also developed a more lightweight browser for unpaired DNA-RNAseq datasets, where input WGS and RNAseq datasets are not derived from the same donor subjects (example browser: https://yanwang271.shinyapps.io/splipath_browser_unpaired_analysis/). The SpliPath R package can be installed from https://github.com/KennaLab/SpliPath. The make_sashimi_like_plot and make_gene_wise_plot function in SpliPath were adapted from make_differential_splicing_plot function in LeafCutter^39^. The analyses in this study were performed using R 3.6.

### *KIF5A* minigene reporter assay

The *KIF5A* minigene reporter construct (Figure. S7) was based on the design described by Cheung et al ^20^. Vector assembly was performed using the Gibson Assembly kit from NEB (Ca. No. E5510S, New England BioLabs)^44^ and primers 1-12 (Table. S7). Sequences were obtained from the plasmids pGEMT-PT2A-GFP-Tdtomato-iRFP670 for the bacteria backbone^45^ and an in-house plasmid containing the EF1alpha-mScarlet and polyA tail sequences. The *KIF5A* minigene sequence was extracted from genomic DNA derived from SH-SY5Y cells. ALS patient and control mutations (Table. S8) were introduced using the site directed mutagenesis kit from NEB and primers 13-21 (Table. S7) (Ca. No. E0554S, New England BioLabs)^46^. Construct sequences were validated by Sanger sequencing. SH-SY5Y cells were cultured in a 24 well plate at a density of 100.000 cells and 1 ml of medium (DMEM with GlutaMAX, 10% FBS and 0.5% Penicillin-Streptomycin). Cells were transfected with 100 ng of plasmid using the Lipofectamine Transfection Reagent (Ca. No. L3000001, ThermoFisher) as per manufactur instructions. After 48 hours of incubation, cells were harvested and stained with DAPI to select live cells. Fluorescence was measured on a BD FACSCanto II Flow Cytometer. Gates were set to select living and single cells using untransfected cells. A compensation protocol was set up using cells transfected with in house plasmids containing eGFP or mScarlet only. The data was analyzed using the FlowJo software. mScarlet signals were obtained from eGFP positive cells to ensure transfection. The exon skipping signal was defined as the median mScarlet signal divided over the median mScarlet signal of the wildtype plasmid. One-way ANOVA followed by T tests (adjusted for multiple testing using *Benjamini–Hochberg* false discovery rate method) were performed to compare the exon skipping signals of wildtype insertions to that of negative control, positive control, and target variants insertions.

## Acknowledgements

We are grateful to the patients with ALS and their families allowing donation of tissue for research. This project has received funding from the European Research Council (ERC) under the European Union’s Horizon 2020 research and innovation programme (grant agreement n° 772376 - EScORIAL. The collaboration project is co-funded by the PPP Allowance made available by Health∼Holland, Top Sector Life Sciences & Health, to stimulate public-private partnerships. K.K. is supported by grants from the Dutch Research Council (grant no. ZonMW-VIDI 91719350) and the ALS Foundation Netherlands. K.K., E.A., J.V. are supported by ALS Stichting (grant “ ALS Tissue Bank – NL ”). All NYGC ALS Consortium activities are supported by the ALS Association (ALSA, 19-SI-459) and the TOW Foundation.

## Consortia

### Project MinE ALS sequencing consortium

Philip van Damme, Philippe Corcia, Philippe Couratier, Patrick Vourc’h, Orla Hardiman, Russell McLaughin, Marc Gotkine, Vivian Drory, Nicola Ticozzi, Vincenzo Silani, Jan H. Veldink, Leonard H. van den Berg, Mamede de Carvalho, Jesus S. Mora Pardina, Monica Povedano, Peter M. Andersen, Markus Weber, Nazli A. Başak, Ammar l-Chalabi, Chris Shaw, Pamela J. Shaw, Karen E. Morrison, John E. Landers, Jonathan D. Glass, Clifton L. Dalgard

### NYGC ALS consortium

Hemali Phatnani, Justin Kwan, Dhruv Sareen, James R. Broach, Zachary Simmons, Ximena Arcila-Londono, Edward B. Lee, Vivianna M. Van Deerlin, Neil A. Shneider, Ernest Fraenkel, Lyle W. Ostrow, Frank Baas, Noah Zaitlen, James D. Berry, Andrea Malaspina, Pietro Fratta, Gregory A. Cox, Leslie M. Thompson, Steve Finkbeiner, Efthimios Dardiotis, Timothy M. Miller, Siddharthan Chandran, Suvankar Pal, Eran Hornstein, Daniel J. MacGowan, Terry Heiman-Patterson, Molly G. Hammell, Nikolaos. A. Patsopoulos, Oleg Butovsky, Joshua Dubnau, Avindra Nath, Robert Bowser, Matthew Harms, Eleonora Aronica, Mary Poss, Jennifer Phillips-Cremins, John Crary, Nazem Atassi, Dale J. Lange, Darius J. Adams, Leonidas Stefanis, Marc Gotkine, Robert H. Baloh, Suma Babu, Towfique Raj, Sabrina Paganoni, Ophir Shalem, Colin Smith, Bin Zhang, Brent Harris, Iris Broce, Vivian Drory, John Ravits, Corey McMillan, Vilas Menon, Lani Wu, Steven Altschuler, Yossef Lerner, Rita Sattler, Kendall Van Keuren-Jensen, Orit Rozenblatt-Rosen, Kerstin Lindblad-Toh, Katharine Nicholson, Peter Gregersen, Jeong-Ho Lee M.D, Maze Therapeutics, Bristol-Myers Squibb, Sulev Koks, Stephen Muljo, Bryan J. Traynor, Pfizer, Regeneron, Insitro

