## Supplementary_figures for "Functional clustering of splice-altering variants in whole genome sequencing data reveals hidden heritability in rare variant disorder"

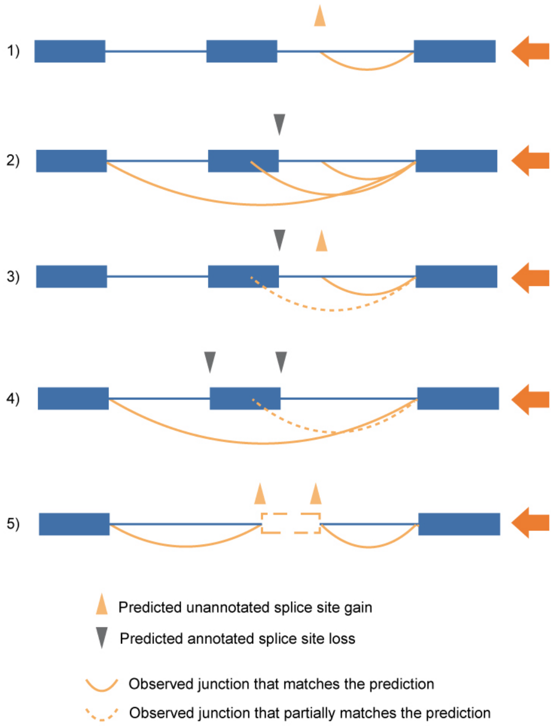


| SpliceAI Prediction | Criteria for “Matched” Junction in RNAseq |
|---|---|
| Unannotated splice site gain | Unannotated junction that uses the predicted splice site. |
| Loss of annotated splice site | Unannotated junction that does use the predicted lost site*. |
| Loss of annotated splice site & gain of unannotated splce site | Unannotated junction that uses the predicted unannotated splice site, and unannotated junction that does use the predicted lost site*. |
| Loss of two annotated splice sites | Unannotated junction that skips both of the predicted lost sites, and unannotated junction that skips one of the predicted lost site*. |
| Gain of two unannotated splice sites | Unannotated junction that uses the predicted unannotated splice site. |

*The lost splice site can be replaced by an alternative annotated site or an unannotated site.

**Figure S1. Evaluating concordance of unannotated junction definitions inferred using SpliceAI predictions and RNAseq seq based split-read analyses.** The orange arrows refer to the predicted unannotated splice sites gain (delta scores ≥ 0.2) and the dark grey arrows refer to the predicted annotated splice sites loss (delta scores ≥ 0.2). The solid orange curves refer to splice junctions that fully match the SpliceAI[1] prediction, and the dashed curves refer to splice junctions that partially match the SpliceAI prediction. The table specifies criteria for SpliceAI prediction “match” the observed junction in RNAseq. When SpliceAI predicts multiple splice sites gain/loss, the junctions match prediction of at least one but not all splice sites gain/loss is considered as “partial match”.

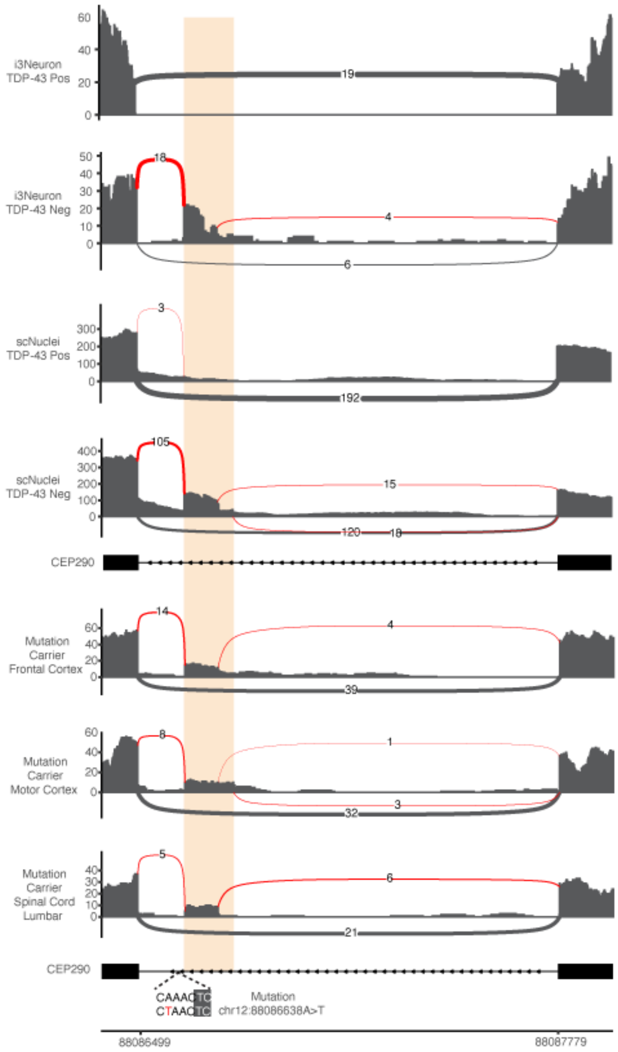


**Figure S2.** ***CEP290*** **cryptic exon inclusion events were observed consistently across tissue types within splice-altering variants carriers.** *CEP290* cryptic exons were observed consistently across cortex and spinal cord samples of chr12:88086638A>T carrier, respectively.

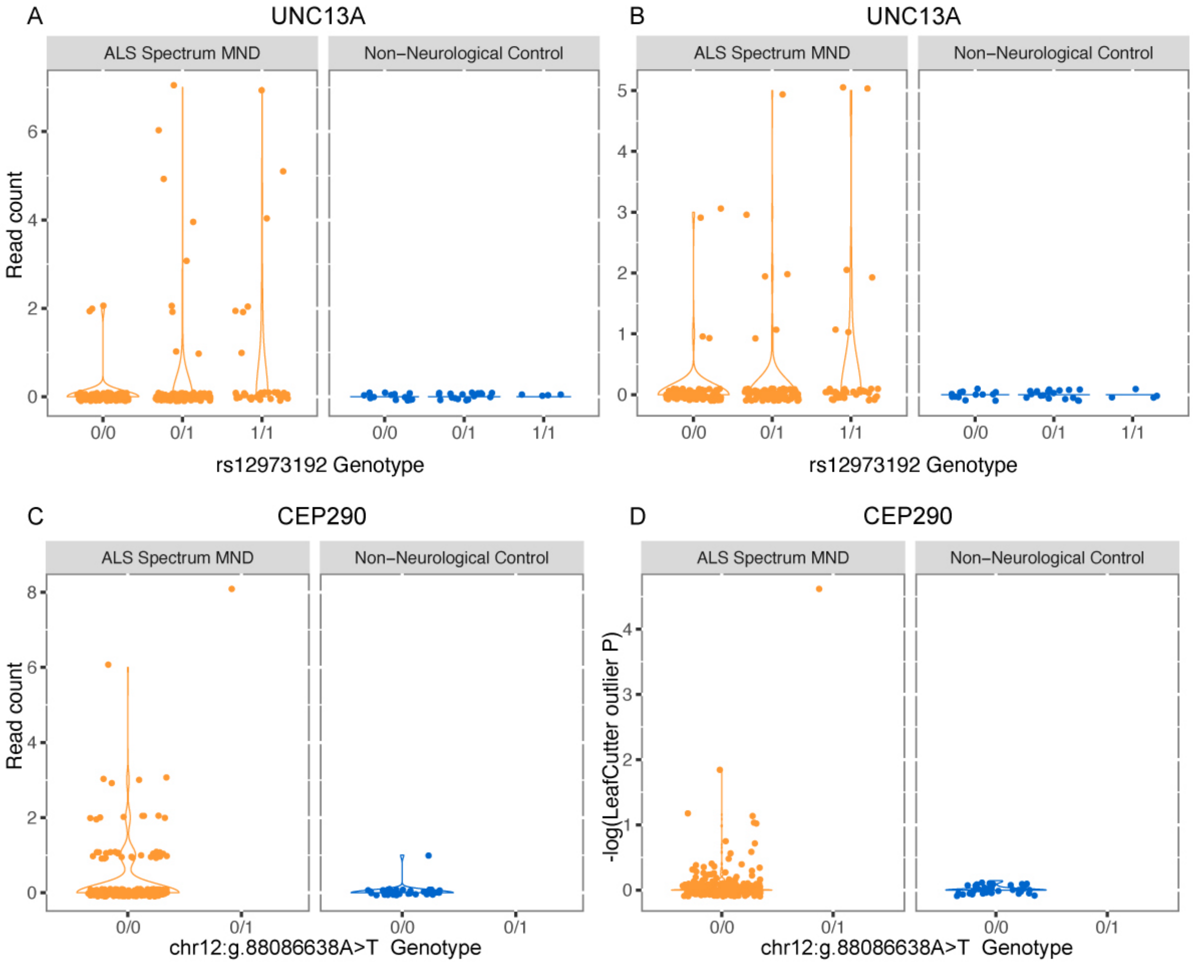


**Figure S3. Junction expression level in tissues of *CEP290* csQTL and *UNC13A* CE modifier carriers.** The number of reads mapped to junction A) chr19:17641556-17642413 and B) chr19:17642541-17642844 in *UNC13A* versus the genotype of CE modifier variant (rs12973192) in NYGC motor cortex samples. The expression level of *CEP290* junction chr12:88086498-88086641, represented by C) the number of reads mapped to the junction and D) the splicing outlier P values, versus the genotype of nominated csQTL (chr12:g.88086638A>T) in NYGC motor cortex samples.

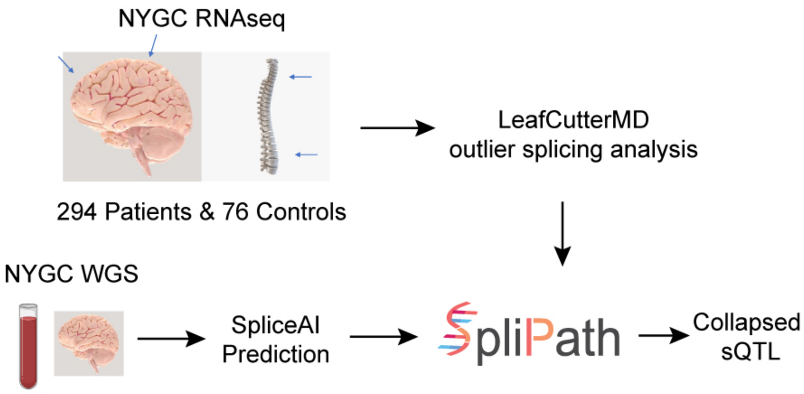


**Figure S4. SpliPath nominates csQTL from paired transcriptomic and genomic data.** In this study, paired RNAseq and WGS data of 294 patients and 76 controls from New York Genome Center ALS consortia are used. The RNAseq samples are from four tissues: motor cortex, frontal cortex, spinal cord cervical and lumbar. The samples of each tissue are analysed by LeafCutterMD to identify outlier splicing junctions. The outlier junctions are then matched with SpliceAI predicted splice-altering variants occurred in the same individual by SpliPath (details in Figure S1). We restricted our study to rare DNA variants, which we defined as those exhibiting a minor allele frequency under 0.1%, or those exhibiting a homozygous carrier frequency under 0.1% in the gnomAD (v3.1.2)[2] non-neurological disease group. Rare variants whose predicted splice-altering effect are observed in RNAseq data are nominated as csQTL.

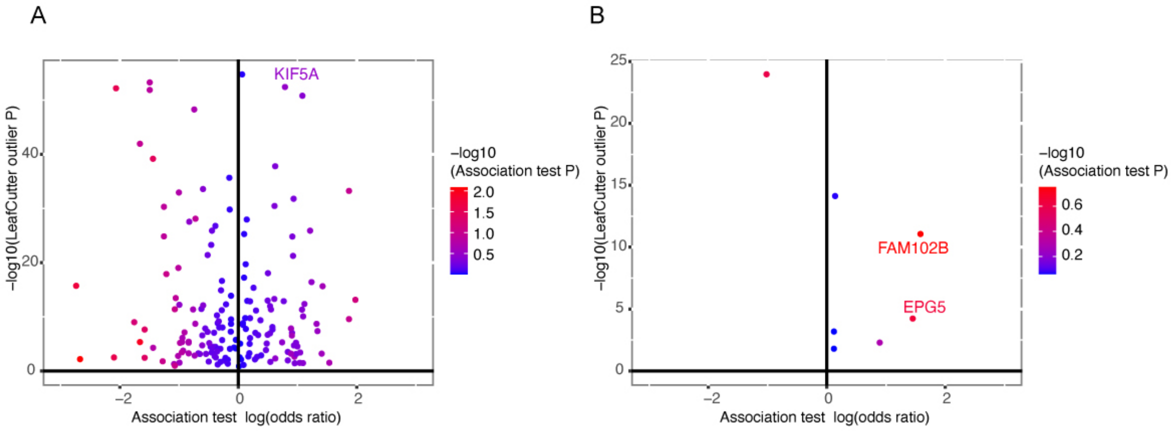


**Figure S5. CsQTL association tests using both allelic and recessive models**. Association tests with A) allelic model and B) recessive model are performed for csQTL annotated by SpliPath using different reference junctions and SpliceAI DNA variant annotations. Each dot shows a junction in NYGC dataset with a outlier splicing P value (Y axis) and the association test odds ratio on log scale (X axis) of the csQTL in Project MinE WGS. Only single nucleotide variants are included in the association tests. The dots are colored by the association test P value.

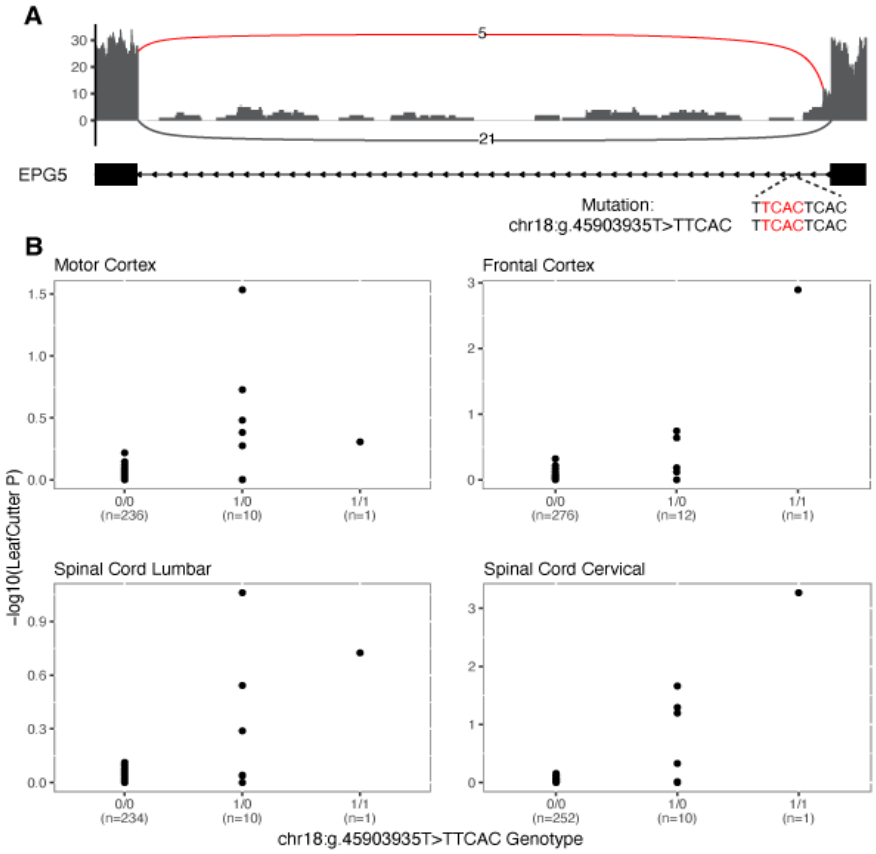


**Figure S6. The *EPG5* unannotated junction express more in tissues of the csQTL carriers**. A) shows the unannotated junction with exon elongation event caused by the variant chr18:g.45903935T>TTCAC in patient's frontal cortex sample. B) are the unannotated junction expression level in donors' tissues harbor different genotype of the csQTL. X axes are the genotype and the number of carriers. Y axes are the expression level of the junction chr18:45901168-45903939, in -log10 transformed LeafCutterMD[3] outlier P value. The junction expression in four tissues, motor cortex, frontal cortex, lumbar and cervical are shown.

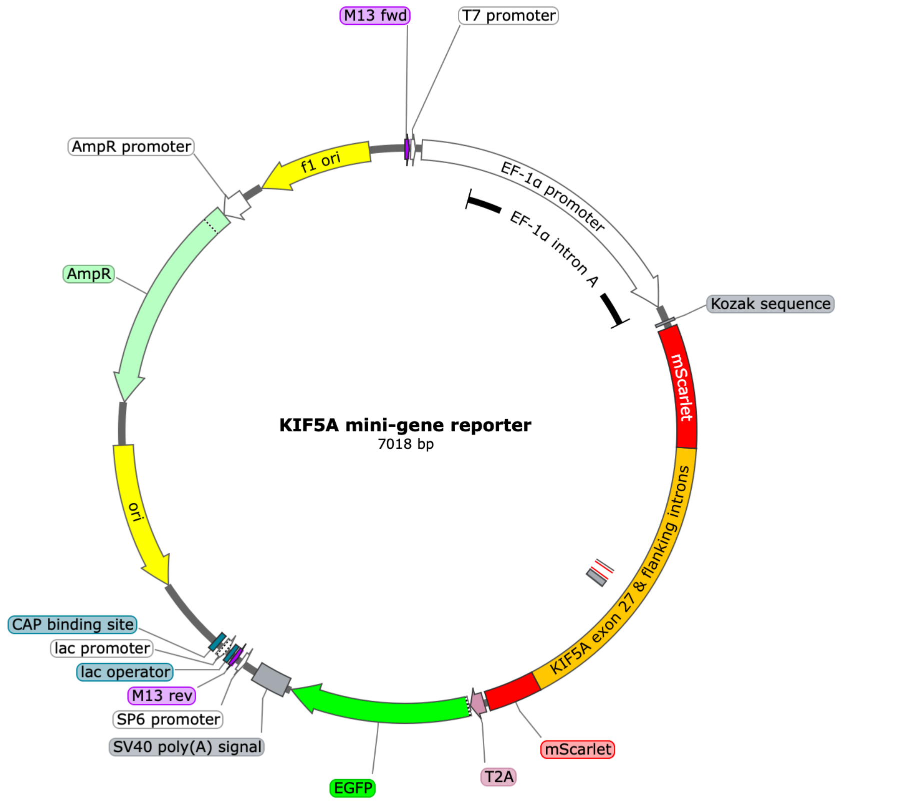


**Figure S7. Design of plasmid for *KIF5A* minigene reporter assay**
